# Context-Dependent Effects of Midkine on Plexiform Neurofibroma Growth and Drug Response in 3D Coculture Models

**DOI:** 10.1101/2025.10.20.683475

**Authors:** Ahmad Zunnu Rain, Harini G. Sundararaghavan, Kyungmin Ji

## Abstract

Midkine (MDK) is a heparin-binding growth factor that promotes tumor growth in many cancers and may contribute to type I neurofibromatosis (NF1) tumor by stimulating Schwann cell proliferation and supporting neurofibroma growth. Here, we investigated the role of midkine in plexiform neurofibroma with NF1 (pNF1) using both monoculture and neuron–tumor coculture system with biomimetic neuronal axons engineered to release midkine in 3D. In monoculture, midkine enhanced the growth of pNF1 tumor cells, consistent with its tumor-promoting role in several cancers. In contrast, in 3D coculture, midkine did not increase tumor growth and failed to alter response to selumetinib, the FDA-approved treatment for pNF1. These findings suggest that while midkine promotes tumor growth, its effect may be buffered or neutralized in a neuron-associated microenvironment, underscoring the importance of physiologically relevant models for evaluating therapeutic targets in NF1.

## Introduction

Neurofibromatosis type 1 (NF1) is a common inherited tumor predisposition syndrome caused by mutations in the *Nf1* gene, which encodes the RAS regulator neurofibromin. Loss of *Nf1* function results in hyperactive RAS/ mitogen-activated protein kinase (MAPK) signaling [1,2] and the development of a wide spectrum of benign and malignant tumors, including plexiform neurofibromas (pNFs), cutaneous neurofibromas, and malignant peripheral nerve sheath tumors (MPNSTs) [3]. While tumor initiation is driven by Schwann cell (SC) lineage cells with *Nf1* loss [1,2], tumor growth and progression are strongly affected by signals from the tumor microenvironment (TME), including neurons, fibroblasts, mast cells, immune cells, and endothelial cells [4,5].

Midkine, a heparin-binding growth factor, has been widely studied as a pro-tumorigenic mediator in cancer biology. In many tumor types, midkine promotes proliferation, survival, angiogenesis, invasion, and metastasis (review [6-8]). In addition to enhancing tumor growth, midkine contributes to therapeutic resistance [9], where it has been shown to attenuate drug efficacy through activation of survival signaling, modulation of apoptosis, and remodeling of the tumor microenvironment. For example, elevated midkine expression is associated with poor prognosis in hepatocellular carcinoma [10], lung cancer [7], breast cancer [11,12], and glioblastoma [13], where it drives tumor expansion and resistance to chemotherapy or targeted therapy [6]. These findings underscore midkine as a multifunctional mediator of cancer progression and treatment resistance.

In NF1, midkine has emerged as a potential regulator of tumor progression. Elevated serum midkine levels have been reported in patients with NF1[14,15], and increased expression is detected in SCs, endothelial cells, and *Nf1*-deficient SCs in mouse models [15,16]. Functionally, midkine promotes proliferation of SCs and endothelial cells [14] and modulates the TME through immune activation. Neuron-derived midkine activates T cells to induce chemokine (CCL) production in pNF1-optic pathway gliomas (OPG) models [17], and in *Nf1*-mutant neurons, midkine stimulates naïve CD8^+^ T cells to produce CCL4, which drives microglial CCL5 secretion essential for low-grade glioma growth [17] .Collectively, these findings implicate midkine as a multifaceted contributor to NF1 tumor biology through effects on proliferation, angiogenesis, and immune signaling, although its role in therapeutic response remains unclear.

Given midkine’s established role in tumor progression and drug resistance in other cancers [6-8], and its suggested activity in NF1-related tumors, it is plausible that midkine may regulate pNF1 tumor growth and therapeutic sensitivity. Yet, most studies of midkine have been performed in monoculture systems [18,19] or in simplified models that only partially incorporate stromal influences [20,21], which do not capture the complex cellular interactions present in pNF1. To address this gap, we developed a biomimetic neuronal axon system capable of secreting midkine and combined it with a 3D coculture of pNF1 tumor cells. This platform enabled us to examine whether midkine affects pNF1 tumor growth and regulates response to selumetinib, an FDA-approved treatment for pNF1.

## Materials and Methods

### Reagents

Reconstituted basement membrane (rBM; Cultrex™ reduced growth factor basement membrane extract) and type I Collagen (rat tail, high concentration) were purchased from Bio-Techne (Minneapolis, MN, USA) and Corning (Corning, NY, USA), respectively. Phenol red-free Dulbecco’s Modified Eagle Medium (DMEM) and MycoZap™ Plus-CL were purchased from Lonza (Basel, Switzerland). Fetal bovine serum (FBS) was obtained from Cytiva (Marlborough, MA). Selumetinib (AZD6244) was purchased from Selleckchem (Houston, TX, USA). LIVE/DEAD™ Viability/Cytotoxicity kit was purchased from ThermoFisher Scientific (Waltham, MA, USA). Poly(ethylene oxide (PEO) and Poly(D,L-lactide-co-glycolide) (PLGA, 75:25) were purchased from Sigma-Aldrich (St. Louis, MO, USA). Irgacure 2959 (I2959) was purchased from BASF Corporation (Vandalia, IL, USA). Human Midkine and Human Midkine Standard ABTS ELISA Development Kit was purchased from PeproTech (Cranbury, NJ, USA).

### Generation of Biomimetic Neuronal Axons Releasing Midkine

PLGA microsphere preparation and loading: Poly(lactic-co-glycolic acid) (PLGA, 75:25) microspheres were prepared using a standard water/oil/water (w/o/w) double emulsion method [22]. Briefly, 0.300 g PLGA was dissolved in 3 mL dichloromethane (DCM) to generate the organic phase. Human midkine (200 μL of 100 ng/mL) was added to this solution to form the internal aqueous phase. The mixture was sonicated using a Vibra Cell ultrasonic processor to create the primary water-in-oil (w/o) emulsion. This emulsion was immediately dispersed into 40 mL of 0.5% (w/v) aqueous poly(vinyl alcohol) (PVA) and vortexed for 30 s to produce the secondary oil-in-water (o/w) emulsion. The emulsion was stirred for 2 h to allow PLGA solidification and DCM evaporation. Microspheres were washed twice with deionized (DI) water at 1,500 rpm to remove residual solvent, then freeze-dried and stored at –20 °C until use.

Electrospinning: Methacrylated hyaluronic acid (HA; 0.08 g) was mixed with 1.6 g of 4% polyethylene oxide (PEO) to prepare the base spin solution. Irgacure 2959 (0.4 g) was added and vortexed to ensure homogeneity, followed by sonication for 15 min. The solution was stored at 4 °C overnight. Human midkine-loaded PLGA microspheres (0.030 g, freeze-dried) and 300 μL DI water were then incorporated into the spin solution and mixed on an orbital shaker. The final solution was loaded into a 3 mL syringe for electrospinning onto coverslips using the following parameters: flow rate, 1.1 mL/h; voltage, 27 kV; mandrel rotation, 1,400 rpm; and ambient humidity, <30%. After electrospinning, the fibers were UV-crosslinked to ensure stability and stored in a desiccant chamber until further use.

Human Midkine Release and ELISA: Midkine release was evaluated by placing electrospun coverslips into the wells of a 12 well plate. Each well was filled with 500 µL of sterile 1× PBS to initiate release of Human Midkine from the microspheres. At predetermined time intervals (30 min, 2 h, 4 h, 8 h, 12 h, 24 h, 48 h, 72 h, 96 h, and 120 h), the PBS from each well was collected and replaced with fresh PBS to continue the release process. After 120 h of sequential collection, the samples were analyzed by ELISA to determine Midkine concentration. Briefly, ELISA plates were coated with a diluted capture antibody and incubated overnight at room temperature. Wells were then washed, and 300 µL of blocking buffer was added to each well, followed by incubation for 1 h at room temperature. After washing, standards and collected samples were added to the wells and incubated for 2 h at room temperature. Subsequently, 100 µL of detection antibody was added to each well and incubated for 2 h. After washing, 5.5 µL of avidin– HRP was added per well and incubated for 30 min at room temperature. Following additional washes, substrate solution was added to each well. Absorbance was measured at 405 nm with wavelength correction at 650 nm. Midkine concentrations were calculated using the standard curve.

### Cell lines and cell maintenance

Two immortalized human plexiform neurofibroma cell lines (*Nf1*^*-/-*^; hereafter called pNF1 tumor cells) described in [23] were used: ipNF95.11b C and ipNF05.5 obtained from ATCC. All cell lines were maintained as monolayers at 37°C, 5% CO_2_ in DMEM/high glucose supplemented with 10% FBS. Routine screening confirmed the absence of mycoplasma contamination.

### Three-dimensional (3D)/4D (3D in real-time) culture models

Three-dimensional (3D) cultures were established either in patented microfluidic culture devices called TAME (tissue architecture and microenvironment engineering; Patent#, US10227556B2) chips[24] as previously published[4,25]. Fabrication of TAME chips, including separate and linked well designs[24], and protocols for 3D monocultures have been described previously [26]. Cells were seeded on a reconstituted basement membrane (rBM; Cultrex™ and collagen I mixture) on glass coverslip in the TAME chips [27]. Cultures were overlaid with DMEM supplemented with 2.5% FBS and 2% rBM and maintained for 5-6 days. There are four experimental groups for 3D culture models (See Fig. 2): 1) rBM ± 40 ng/ml of midkine on glass coverslip for pNF1 monocultures; 2) rBM on biomimetic neuronal axons with empty PLGA or midkine-releasing PLGA on glass coverslip for pNF1:biomimetic neuronal axon cocultures. Cells in midkine-treated groups are exposed to midkine in both rBM and culture media. Prior to live-cell 3D imaging, nuclei were stained with Hoechst 33342 to assess cell integrity and morphology, and total live-cell numbers were quantified using viability markers such as Calcein-AM.

### Image acquisition for quantitative analysis in 3D

Optical sections spanning the full depth of the 3D structures were acquired from 4 contiguous fields (2 × 2) using an inverted Leica Stellaris 5 system (Leica Microsystems, Deerfield, IL, USA) equipped with a Tokai Hit stage-top incubator to maintain physiological conditions. Image stacks were processed and reconstructed in 3D using Volocity software (Version 7.0.0; PerkinElmer, Waltham, MA, USA) to visualize and quantify live/dead cell distributions. Spatial orientation within reconstructed volumes is indicated by x (green), y (red), and z (blue) arrows in the lower corner of each image.

### Statistical analysis

Data are presented as bar graphs, representing the mean ± standard deviation (SD). The number of replicates is provided in each corresponding figure legend. Statistical significance was assessed using either one-way ANOVA followed by Tukey’s post-hoc test for multiple group comparisons, or Student’s t-test for comparisons between two groups. A *p*-value ≤ 0.05 was considered statistically significant throughout the study.

## Results

### Establishment of biomimetic neuronal axons and pNF1-neuronal axon 3D cocultures

To investigate the contribution of neuronal midkine to pNF1, we engineered biomimetic neuronal axons incorporating midkine release. Hyaluronic acid–based scaffolds were electrospun to form aligned fiber networks, and midkine-loaded PLGA microspheres were embedded into the constructs. Release kinetics analysis showed that ∼40 ng/mL of midkine was released into rBM over a 120-hour period, consistent with sustained delivery (Fig. 1C). When layered into the 3D culture system, the platform established three distinct compartments: (i) biomimetic neuronal axons (green) at the base, (ii) rBM matrix in the middle, and (iii) pNF1 tumor structures seeded on top (Fig. 1D). This engineered microenvironment provided a physiologically relevant model for examining midkine-dependent effects on tumor growth and drug response.

**Fig. 1.**
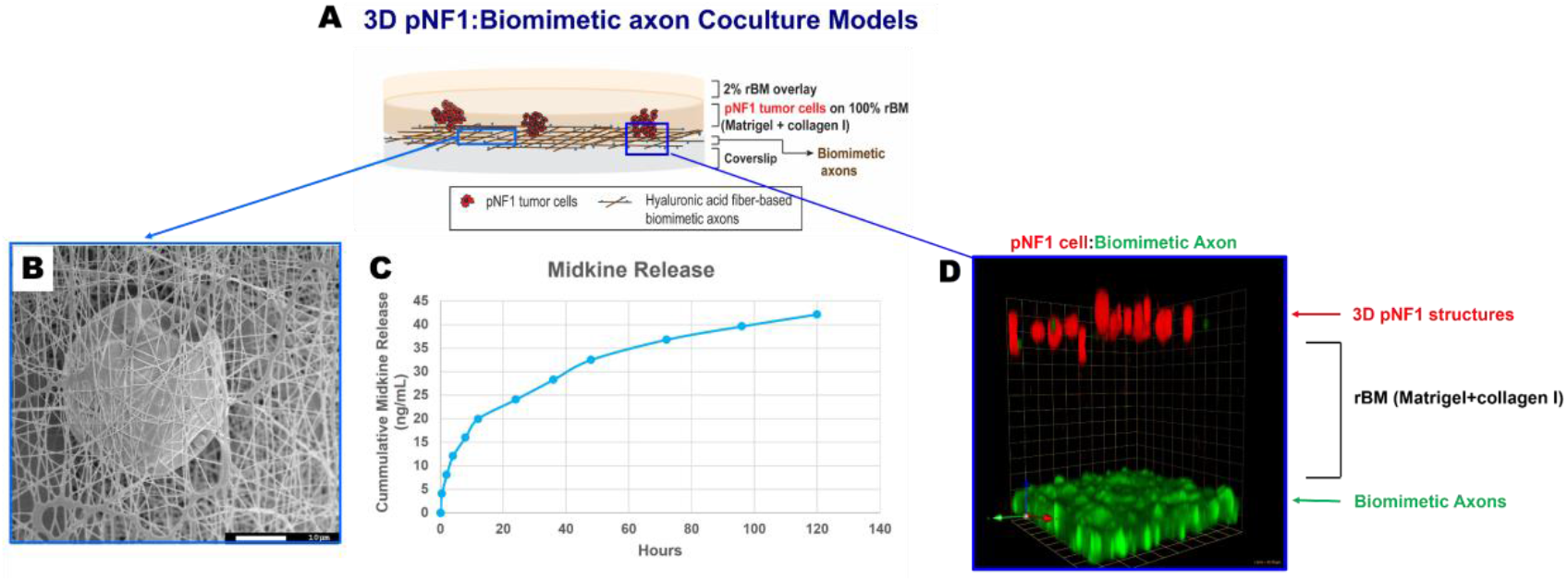
Establishment of 3D coculture of pNF1 tumor cells and biomimetic neuronal axons releasing midkine. (A) Schematic illustration of the 3D coculture system, showing 3D pNF1 structures layered with biomimetic neuronal axons. (B) Scanning electron microscopy (SEM) image of hyaluronic acid (HA) fibers containing PLGA microspheres loaded with midkine (MDK). Scale bar, 10 μm. (C) Cumulative release profile of midkine from PLGA microspheres over 7 days. (D) Side-view 3D reconstruction of 3D pNF1 structures formed by ipNF95.11bC (red) cocultured with biomimetic neuronal axons (green) for 7 days. Grid, 43 μm.

### Midkine increases tumor growth in monoculture but not in coculture with neuronal axons in 3D

We next examined the effect of midkine on 3D pNF1 structures formed by ipNF95.11bC or ipNF05.5 cells grown in the TAME chips. Four conditions were tested: 1) pNF1 monocultures on rBM-coated glass coverslips (control for monocultures; Fig. 2A, top); 2) pNF1 monocultures on rBM containing 40 ng/mL midkine with midkine also added to the culture medium (Fig. 2A, bottom); 3) pNF1:biomimetic neuronal axon cocultures on rBM with empty PLGA microspheres (control for cocultures; Fig. 2B, top); and 4) pNF1:biomimetic neuronal axon cocultures on rBM with midkine-releasing PLGA microspheres (Fig. 2B, bottom). In 3D monocultures, treatment with midkine significantly enhanced tumor growth compared with untreated controls, as quantified by viability assays (Fig. 2A, E). In contrast, in the 3D coculture system containing biomimetic neuronal axons with sustained midkine release, no significant differences in tumor growth were observed relative to controls (Fig. 2B, E). These findings indicate that while midkine promotes pNF1 tumor growth in 3D monocultures, its effect is diminished in a tumor– neuron coculture environment, suggesting that neuronal axons may provide compensatory signals that counteract midkine-driven growth.

**Fig. 2.**
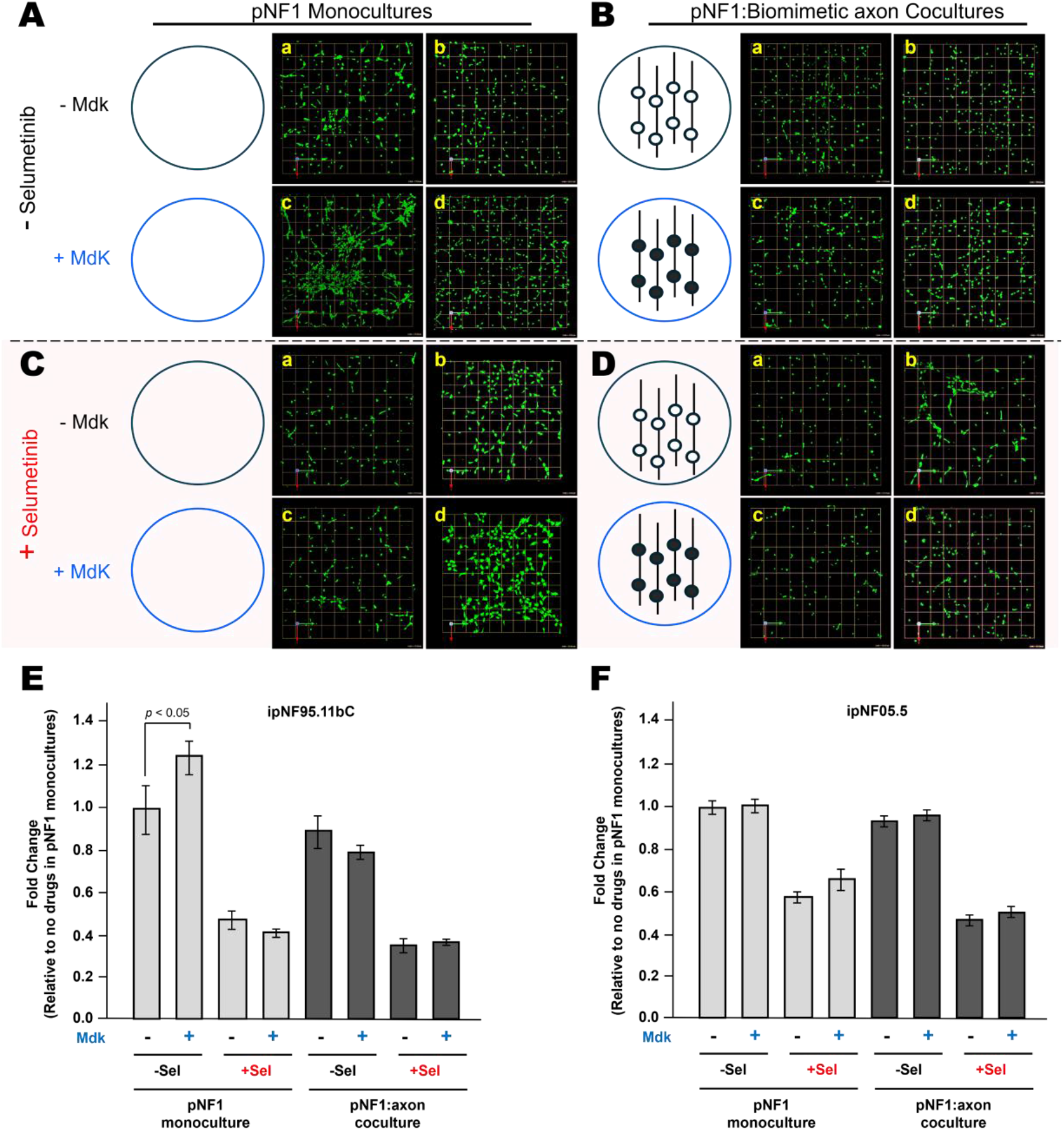
Midkine promotes pNF1 tumor growth in monoculture but not in coculture with biomimetic neuronal axons, and does not alter selumetinib sensitivity. (A–D) Schematic illustration of 3D culture conditions: (A) pNF1 monocultures ± midkine, (B) pNF1:biomimetic neuronal axon cocultures ± midkine, (C) monocultures ± selumetinib, and (D) cocultures ± selumetinib. 3D tumor structures derived from ipNF95.11bC (middle panels; a and c) or ipNF05.5 cells (right panels; b and d) were embedded in rBM with or without midkine and treated ± selumetinib for 5 days. Tumor structures were stained with calcein AM (green, live-cell marker). Images are tiled from 4 contiguous fields; grid size = 174 μm. (E) Quantification of viable pNF1 cells using Volocity software. Data represent mean ± SEM (n = 3).

### Midkine does not alter selumetinib response in 3D pNF1 tumor structures

To assess whether midkine affects therapeutic sensitivity, 3D pNF1 tumor structures were treated with selumetinib for 5 days in the presence or absence of midkine. In monocultures, selumetinib alone reduced the number of viable tumor structures (Fig. 2C, top). Co-treatment with midkine did not alter the inhibitory effect of selumetinib, as shown by viability analyses (Fig. 2C, D). Similarly, in the biomimetic coculture system, selumetinib suppressed pNF1 tumor growth to a comparable extent regardless of midkine exposure (Fig. 3C). These findings indicate that midkine does not confer resistance to selumetinib in pNF1, at least within the 3D *in vitro* models tested. Overall, our results show that midkine promotes tumor growth in monoculture but not in neuronal coculture, and it does not affect selumetinib response, underscoring the context-dependent role of midkine in pNF1.

## Discussion

This study demonstrates that the effect of midkine on pNF1 tumor cells is strongly dependent on the experimental context. In monoculture, midkine promoted pNF1 tumor cell growth, consistent with prior reports that MDK functions as a mitogen in Schwann cells [14]. Elevated circulating midkine levels have also been reported in pNF1 patients [14,15], supporting the idea that midkine is biologically active in this tumor type. However, in a 3D coculture system incorporating biomimetic neuronal axons engineered to release midkine, the growth-promoting effect was absent, and no influence on selumetinib response was detected.

Our findings appear to contrast with the broader literature, in which midkine is consistently described as a potent mitogen and driver of therapeutic resistance [9]. Several possible explanations may account for these discrepancies. First, differences in tumor type may be critical. In NF1-OPG, midkine functions within a glial-neuronal niche in which neuronal activity strongly supports glioma initiation and progression [17]. In contrast, pNF1 tumors originate from Schwann cell lineages and are embedded in the TME with fibroblasts, mast cells, and endothelial cells [5]. It is possible that the cellular receptors and downstream signaling pathways for midkine differ between glial and Schwann cell contexts, leading to divergent outcomes.

Second, the experimental model likely shapes the observed effect. Many published studies have relied on two-dimensional cultures [18,19] or *in vivo* systems where stromal, neuronal, and immune contributions overlap[16,21]. Our 3D biomimetic axon system provides a controlled neuronal input but may lack additional stromal and immune cues that are necessary for midkine-driven tumor promotion. For instance, midkine has been shown to stimulate endothelial cell proliferation [14], a function not fully captured in our 3D coculture model. Thus, the absence of growth promotion or drug resistance in our system may reflect the lack of endothelial and immune components that amplify midkine activity *in vivo*.

Third, midkine signaling is likely dose- and context-dependent. *In vivo*, midkine secretion is regulated by neuronal activity in a spatially and temporally dynamic fashion, whereas our biomimetic model provides constitutive midkine release. This difference may alter downstream signaling dynamics and attenuate midkine’s apparent activity.

Finally, tumor stage and microenvironmental maturation could also account for the discrepancies. OPGs often arise in young children, where neuronal and glial circuits are developmentally plastic and may be particularly sensitive to midkine signaling [17]. In contrast, the adult-derived pNF1 tumor cells used here may have different receptor repertoires or adaptive mechanisms that diminish midkine responsiveness [12,28,29], and microenvironmental maturation can alter therapeutic and signaling outcomes [30].

Taken together, these considerations suggest that midkine’s role in NF1 tumor is not uniform across tumor types or model systems. In monocultures, midkine reveals its canonical growth-promoting effects. In coculture systems that more closely mimic the neuronal microenvironment, its activity appears constrained by redundant or competing pathways.

The implications of these results are two-fold. From a mechanistic standpoint, they highlight the complexity of neuron–tumor signaling in NF1 and emphasize that paracrine factors act within a network rather than in isolation. From a translational standpoint, they caution against extrapolating monoculture findings to therapeutic strategies, since factors like midkine may have attenuated roles *in vivo*. Ultimately, our results reinforce the value of physiologically relevant models for identifying actionable therapeutic targets in NF1-associated tumors.

## Patents

The present study is associated with the U.S. Patent US10227556B2.

## Author Contributions

Conceptualization, K.J.; investigation, K.J. and A.Z.R.; writing-original draft, K.J.; writing-review and editing, K.J., A.Z.R., and H.G.S.; supervision, K.J. and H.G. S. All authors have read and agreed to the published version of the manuscript.

## Acknowledgments

We acknowledge the support and contributions of Bonnie F. Sloane, K.J.’s great mentor, who passed away from lung cancer and gave feedback and encouragement on the development of NF1 projects. We thank K.J.’s NF1 mentoring committee, Raymond Mattingly at East Carolina University Brody Medical School (Greenville, NC, USA) and Katherine Gurdziel at Wayne State University (Detroit, MI, USA) for their input and encouragement on the overall NF1 projects through a monthly NF1 meeting. We also thank the Neurofibromatosis Therapeutic Acceleration Program (NTAP) for its funding support in the purchase of immortalized human pNF1 cell lines for this study.

